# Novel algorithms for improved detection and analysis of fluorescent signal fluctuations

**DOI:** 10.1101/2022.08.03.502593

**Authors:** Gebhard Stopper, Laura C. Caudal, Phillip Rieder, Davide Gobbo, Lisa Felix, Katharina Everaerts, Xianshu Bai, Laura Stopper, Christine R. Rose, Anja Scheller, Frank Kirchhoff

**Author notes:** Correspondence to: Prof. Dr. Frank Kirchhoff, Molecular Physiology, CIPMM, University of Saarland, 66421 Homburg, Germany.

## Abstract

Fluorescent dyes and genetically encoded fluorescence indicators (GEFI) are common tools for visualizing concentration changes of specific ions and messenger molecules during intra-as well as intercellular communication. Using advanced imaging technologies, fluorescence indicators are a prerequisite for the analysis of physiological molecular signaling. Automated detection and avnalysis of fluorescence signals requires to overcome several challenges, including correct estimation of fluorescence fluctuations at basal concentrations of messenger molecules, detection and extraction of events themselves as well as proper segmentation of neighboring events. Moreover, event detection algorithms need to be sensitive enough to accurately capture localized and low amplitude events exhibiting a limited spatial extent. Here, we present two algorithms (PBasE and CoRoDe) for accurate baseline estimation of fluorescent detection of messenger molecules and automated detection of fluorescence fluctuations.

**Author summary:** Analyzing molecular signalling is crucial in understanding intra- and intercellular communication. These signals are visualized using fluorescent dyes or genetically encoded fluorescence indicators. In the brain, Ca^2+^ signals of glial cells are essential in deciphering complex regulatory functions in health and disease. Due to signal heterogeneity, detection and analysis are highly challenging. They can be stationary, with low amplitude and localized in cell processes, occur as prominent somatic signals or propagate as waves across cellular networks.

We have developed two algorithms to analyze fluorescence transients, each tackling a specific problem. PBasE performs automatic and adaptive background correction, removing basal fluorescence fluctuations. CoRoDe automatically extracts regions of interest, explicitly including temporal information to obtain a precise segmentation, which is essential for accurate transient extraction. Combined, these algorithms are able to detect regions exhibiting low amplitude transients with small spatial extent as well as large, high amplitude signals. Extracted transients are categorized based on their peak amplitude, allowing detailed analyses by comparing changes of specific properties. In order to make these algorithms accessible, an interactive application, called Msparkles, has been designed.

## Introduction

Analysis of ligand dependent fluorescence fluctuations in the central nervous system is a major step in unravelling complex regulatory functions. Astroglial Ca^2+^ events are considered a key factor in identifying the versatile roles of astrocytes in health and disease (1-4). However, reliable detection, analysis and interpretation of such Ca^2+^ events appear to be a non-trivial task (5-11). One of the challenges in analyzing astroglial Ca^2+^ events lies in the heterogeneous nature of astrocytes themselves, reflected in a complex morphology, diverse interactions with other cell types ranging from vasculature to synapses and the formation of intercellular gap junction coupled syncytia (4, 12, 13). The majority of astroglial Ca^2+^ events occur in fine and highly ramified astroglial processes (14), localized at end feet (15) or at synaptic compartments (7). These so-called microdomain events are elementary signals, serving autonomous functions (16), they are not location specific and can occur at all regions of the complex process network. Occurring in functionally independent cellular sub compartments, they occupy volumes in the sub-µm³ range and cause changes in fluorescence close to noise level (14, 17), rendering these events particularly difficult to characterize, requiring careful detection and removal of fluorescence background (17).

Fluorescence events are classically analyzed using stationary regions of interest (ROIs). Recently, new approaches utilizing so-called dynamic events (9, 17, 18) to extract and analyze non-stationary fluorescence events have been proposed. The analysis of dynamic events is an important extension and can reveal new insights. However, most fluorescence events, including the majority of astroglial Ca^2+^ events, are stationary (9), exhibiting only small to no changes in event morphology and location. For this reason, classical ROI analysis remains a valid and powerful tool to quantify fluorescence events and will be focused in this work. Manually outlining fluorescence events is a cumbersome task, experience-dependent and subject to bias. Automatic event detection on the other hand often requires tedious parameter tuning and trade-offs between detection accuracy and sensitivity.

Here, we present two novel algorithms, called PBasE (**P**olynomial **Bas**eline **E**stimation) and CoRoDe (**Co**rrelation-based **RO**I **De**tection). Both algorithms are designed to work independent of specific fluorophores and cell types. The aim of PBasE is to accurately estimate basal fluorescence levels (*F*_0_) while simultaneously preserving fluorescent transients originating from phasic signaling. While PBasE builds upon the idea of penalizing potential fluorescence signals for the purpose of background estimation (17), an additional design goal is the ability to preserve slow, and long lasting fluorescence elevations. Building upon the background corrected dataset ((*F* − *F*_0_)/*F*_0_ = Δ*F*/*F*_0_), CoRoDe is aimed at extracting and segmenting fluorescently active regions by explicitly leveraging temporal information. These algorithms are made accessible via a graphical and interactive analysis application, developed in MATLAB (19), called MSparkles (Suppl. Figure 1 A). MSparkles itself is developed to aid the analysis process by providing direct visual feedback to the user, allowing to explore datasets and quickly optimizing analysis parameters.

The capabilities of PBasE, CoRoDe as well as MSparkles are demonstrated by analyzing differences in astroglial Ca^2+^ events, recorded in the somatosensory cortex of awake and anesthetized mice. These differences have been studied before (17, 20, 21), serving as reference to our results. Next, the results obtained by MSparkles are compared to those obtained with three other Ca^2+^ analysis tools, CHIPS (22), CaSCaDe (7) and AQuA (9). CHIPS provides a collection of various classes and algorithms for specific events, usable with little programming skills. CaSCaDe combines an activity-based ROI detection with a machine learning approach for event extraction. AQuA is based on machine learning principles and uses a combination of thresholding and probabilistic modeling to extract dynamic events. Finally, the use of MSparkles in analyzing fluorescence events is tested in two different experimental situations - Ca^2+^ signals obtained in GCaMP3 reporter mice, as well as neuronal Na^+^ signals visualized using SBFI-AM.

## Results

We developed two novel algorithms to improve the automated detection of fluorescence events and their subsequent analysis. These algorithms were implemented in MATLAB and integrated into a graphical analysis application, called MSparkles. The algorithms were tested by analyzing Ca^2+^ events, visualized using GCaMP3, as well as Na^+^ changes, visualized using SBFI-AM in astrocytes and neurons, respectively. In addition, the results obtained with MSparkles were compared to other analysis applications. The datasets exhibited a broad range of fluorescence events, ranging from somatic events to events in the gliapil, from dim to bright events, varying spatial extent as well as stationary events, dynamic events as well as global events extending throughout the recorded field of view.

### *F*_0_-estimation

PBasE (**P**olynomial **Bas**eline **E**stimation) is an automated, pixel-based algorithm to estimate fluorescence levels at basal concentrations (*F*_0_) of Ca^2+^and other important messenger molecules (Figure 1 A, B). It operates solely along the temporal axes of a dataset, and is thus equally well suited for datasets containing two or three spatial dimensions. First, the algorithm performs, signal clean-up and simplification. Signal clean-up excludes statistically large values from *F*_0_ estimation, and is implemented in two variants. A temporal mean filter computes the mean value (µ) and the corresponding standard deviation (σ) over a pixel’s entire time-course. Then, all values > *µ* + *nσ* are eliminated, where *n* is a user-definable factor. Alternatively, a Hampel filter (23) can be used, which is the sliding window counterpart to the temporal mean filter. To avoid high frequency oscillations of the fitted polynomial, especially near the beginning and the end of the cleaned signal, signal simplification is performed, using piece-wise constant functions to compute a guidance signal. Therefore, a scale space with *k* scales, each containing 2^*k*^ sections, where *k* ∈ [1.. *m*] is computed. Each constant section is computed within an interval 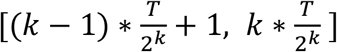, where *T* is the number of recorded time points. The guidance signal can be optimized with respect to local minima, maxima, or minimal error to the cleaned-up signal within the intervals. Finally, a polynomial curve of user-definable degree is fitted to the optimized guidance signal in a least squares sense.

**Figure 1:**
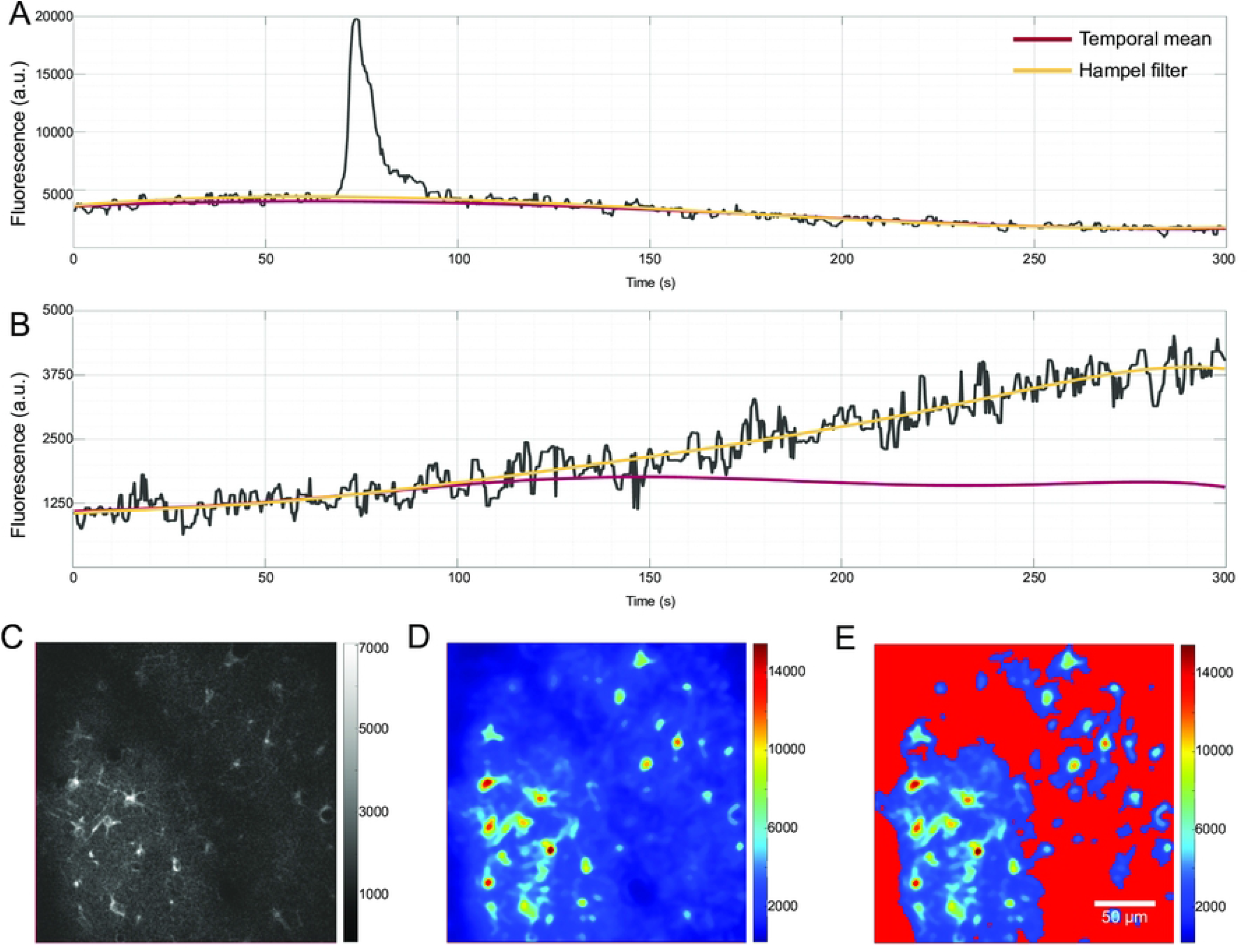
F_0_ estimation by polynomial fitting. From the recorded signal of a single pixel (A, B gray line) potential transients were excluded for baseline estimation either by a Hampel filter or by using the overall temporal mean of a pixel’s signal. A polynomial is then fitted to the ‘clean’ signal in a least-squares sense in order to obtain the estimate of the basal fluorescence level (red, F_0_ estimation based on temporal mean; yellow F_0_ estimation based on Hampel filter). A) Both methods return similar estimates if no long-lasting and slow increases in basal fluorescence are present. B) The temporal mean filter is able to preserve long lasting and slow changes, while the Hampel filter incorporates them into the estimated baseline. C) An individual frame of the original image stack did not reveal fluorescently active regions. D) The fluorescence range projection revealed regions with fluctuations in fluorescence. E) F_0_ masking based on the range projection allows to exclude regions with little or no fluorescent activity from the F_0_ estimation, and effectively prevented the detection of false-positive ROIs and subsequent false transients.

### *F*_0_-masking

Normalizing and detrending a dataset by computing Δ*F*/*F*_0_ may cause artificially amplified artifacts in dark, noisy image regions with no fluorescence activity. Independent of the method used to estimate *F*_0_, values of 1 > *F*_0_ ≥ 0 can occur. This is likely to amplify noise and may lead to the detection of false-positive events, ultimately resulting in the detection of false transients. This problem is solved by *F*_0_-masking, similar to (24). For this purpose, the original stack *F* (Figure 1 C) is used to compute the range projection *R* with *R* = *max*(Δ*F*/*F*_0_ (*p*)) − min (Δ*F*/*F*_0_ (*p*)) (Figure 1 D) along the temporal axes. By applying a user-definable threshold to the range projection, requiring a minimal fluorescence change the *F*_0_ mask (Figure 1 E) is obtained. An initial threshold value is estimated using Otsu’s method, instead of using a fixed percentile as in (24). Pixels covered by the *F*_0_-mask are excluded from the *F*_0_ estimation.

### ROI-detection

Detection and segmentation of events into ROIs is performed by the CoRoDe algorithm (**Co**rrelation-based **Ro**i **De**tection, Figure 2) and is based on the range projection *R*. Pixels excluded by the *F*_0_-mask are zero in *R*. CoRoDe uses simultaneous region growing of local maxima, constrained by a minimally required temporal correlation of neighboring pixels in addition to a range threshold, *t*_*R*_. Local maxima are grown into their neighborhoods until either *t*_*R*_ or the correlation threshold (*t*_*corr*_) is violated. Pixels adjacent to more than one region are marked as boundary pixels. Decreasing *t*_*corr*_ to zero results in region growing being governed by the range threshold, and the detected regions become more similar, but not identical, to those obtained by a watershed segmentation Figure 2 A, bottom).

**Figure 2:**
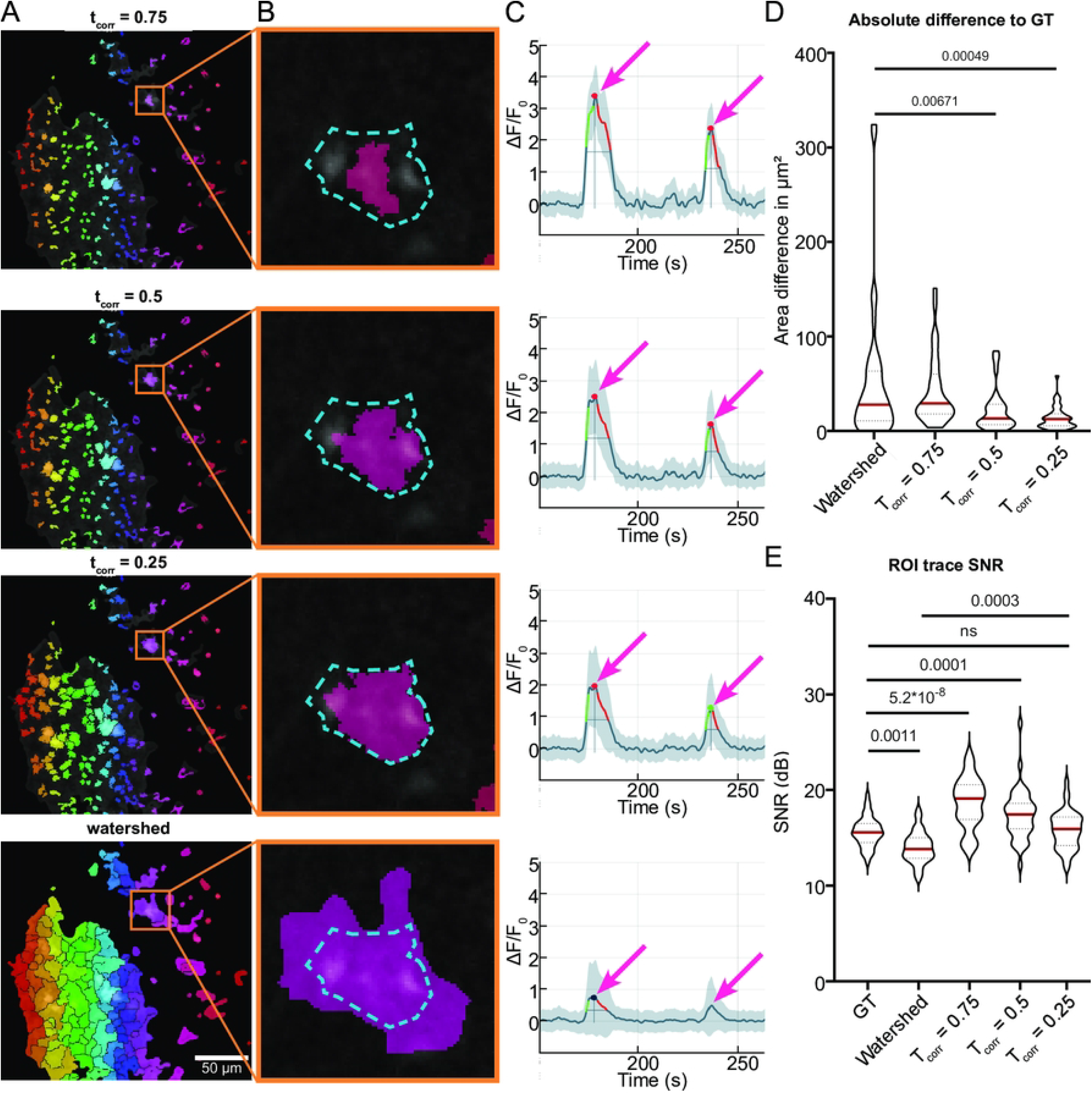
Temporal correlation-based ROI detection. A) ROIs obtained using CoRoDe with different correlation thresholds (t_corr_) as well as watershed transform. Correlation thresholds were set to 0.75, 0.5 and 0.25. B) Direct comparison of automatically extracted ROIs to a manually evaluated ground truth. The obtained ROIs (pink) were contrasted to the manually extracted maximal extent of the fluorescence event (dashed blue line). C) Fluorescence profiles from the highlighted ROIs showed the influence of accurate segmentation on resulting peak amplitudes. Transients of ROIs obtained by watershed segmentation exhibit too small peak amplitudes, while transient peaks of undersized ROIs were too high (arrows). D) ROIs obtained using CoRoDe showed significantly less difference from ground truth, compared to watershed transform. E) Integrated fluorescence profiles from ROIs with appropriate t_corr_ show no difference in SNR, compared to ground truth, whereas profiles from ROIs obtained using a watershed transform showed a significantly reduced SNR.

### Transient analysis and classification

ROI integration, transient extraction and classification are automatically performed on Δ*F*/*F*_0_ in MSparkles. Also, transient amplitudes, durations, rise and decay times (Suppl. Figure 2 A, Suppl. Table 1) as well as inter-transient timings between consecutive transients are automatically determined (Suppl. figure 2 B, Suppl. Table 1). Transient durations can be computed as the full width at half maximum (FWHM), full width at 25% or 10% of the peak amplitude. The latter two can lead to a much more accurate estimation of transient duration, but set higher requirements to signal quality. Optionally, sub-transients can be excluded. Sub-transients are non-maximal peaks of a multi-peak transient, i.e. amplitude peaks occurring during the rise or decay of a stronger peak. This strongly depends on the level at which transient duration is computed.

Extracted transients are automatically classified into user-definable classes, based on their peak amplitude. By default, three pre-defined classification intervals [0.5..1.0), [1.0..1.5) and [1.5.. ∞) are used. Detected ROIs not exceeding the lowest classification threshold (here 0.5 Δ*F*/*F*_0_) at any time are considered false positives and are automatically removed. Classified transients are then used to compute the signal composition, as the relative frequency of transients belonging to a specific class (Figure 3 J, M), revealing changes in signaling behavior.

**Figure 3:**
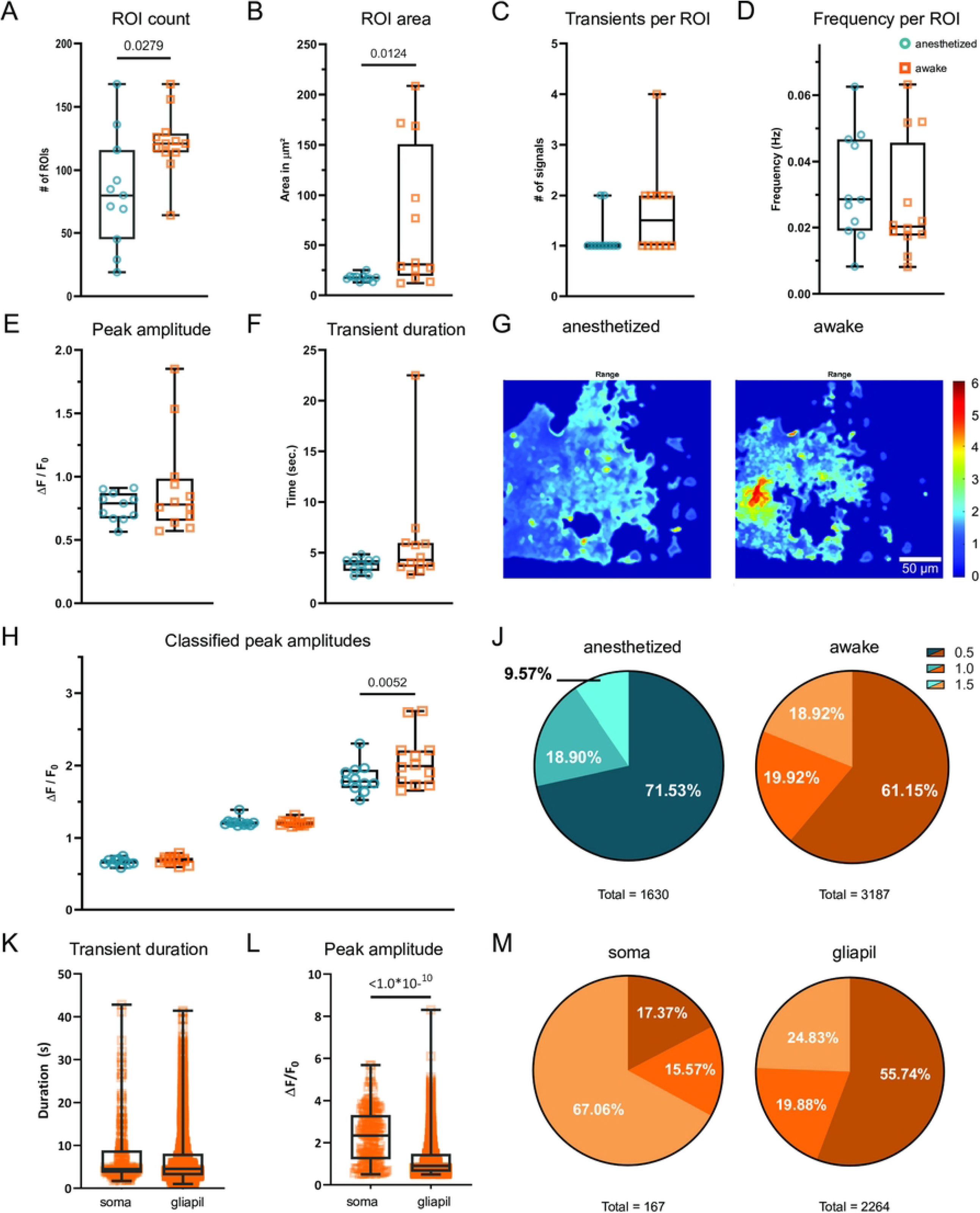
Statistical analysis and transient characterization of Ca^2+^ signals in GCaMP3 mice. The median number of detected ROIs (A), as well as the ROI area (B) increased in awake animals. The median number of transients per ROI also doubled in awake animals expressing GCaMP3 (C). Per-ROI transient frequencies did not change (D). The overall median transient peak in awake animals did not change (E). Overall median transient duration showed no change in awake GCaMP3 animals (F). The range projection of ΔF/F_0_ indicates the amplitude and extent of fluorescence fluctuations (G). Individual classes (H, J) show not only an increase in median amplitude above 1.5 ΔF/F_0_ (H) but also a relative increase (signal composition) in strong transients during wakefulness (J). Differential analysis of somatic transients and transients occurring in the gliapil (K, L, M) shows similar median durations (K). Somatic transients exhibit not only a higher median peak amplitude (L), but also occur mostly as high amplitude transients (M), compared to transients in the gliapil.

### Synchronicity analysis

Astroglial networks can exhibit highly synchronized signaling behavior (13). To detect and analyze temporally correlated events, a synchronicity index, ranging from 0 to 1, as the relative frequency of simultaneously active ROIs per timepoint is automatically computed (Figure 4 A, bottom). A ROI is considered active during the period between the computed start and end time of an identified transient peak. Peaks in the synchronicity index and their respective duration (FWHM) are automatically extracted. All ROIs exhibiting a transient peak in Δ*F*/*F*_0_ during the duration of a synchronous event are extracted together with their activation sequence. Activation sequences are determined, based on the starting times of affected transients. This not only allows to identify synchronously active regions, but also the internal activation pattern and spread of a synchronous event.

**Figure 4:**
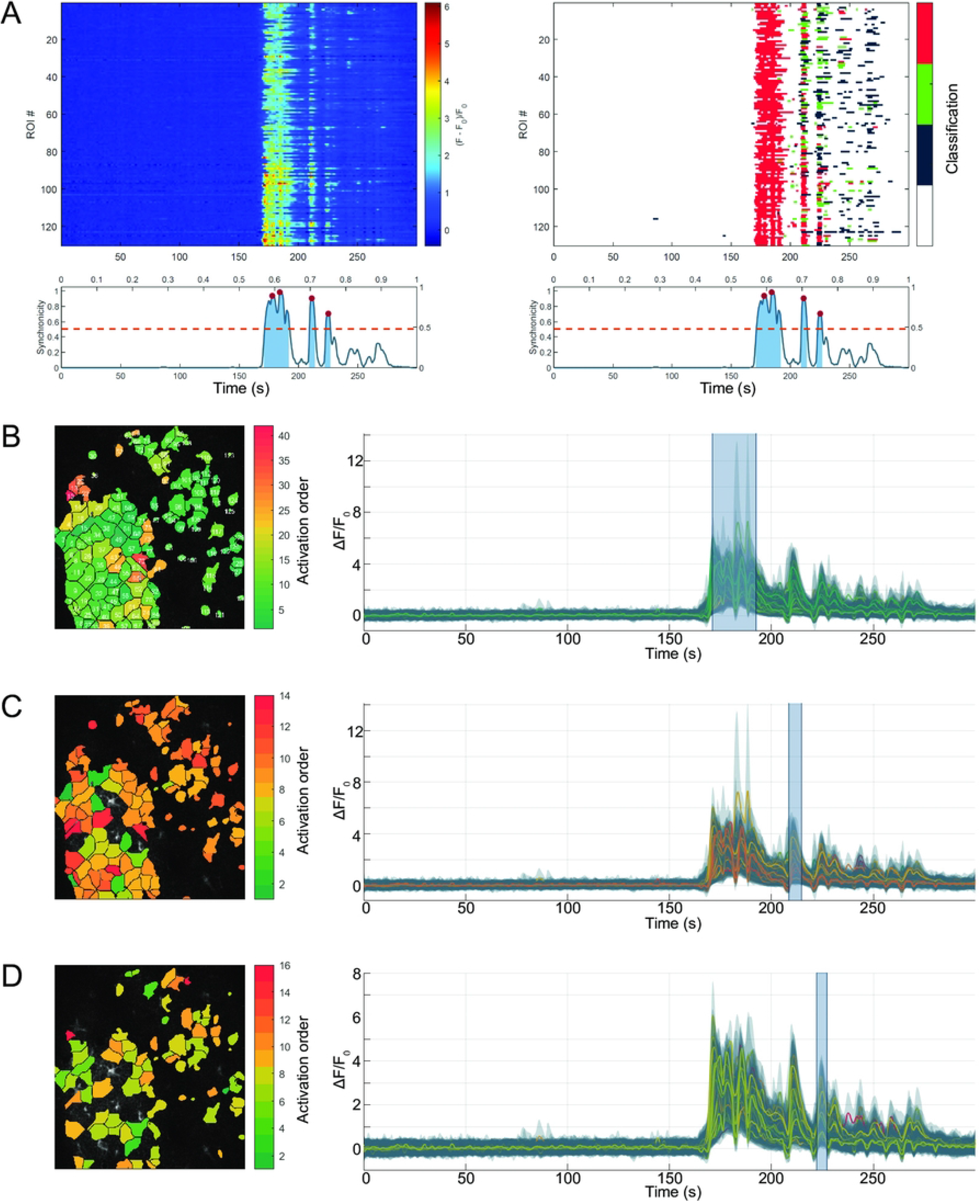
Synchronous events are diverse. Three separate, but consecutive synchronous events in the same recording. Synchronous events are qualitatively assessable via kymographs, showing unclassified fluorescence profiles (A, left) as well as classified transients (A, right). In addition, synchronicity plots (A, beneath kymographs) allow to quantify the relative frequency of synchronous activity, with respect to the number of detected ROIs. Synchronous periods above threshold (red, dashed line) are highlighted (blue areas), and peaks in synchronicity are marked (red circles). Analyzing individual synchronous events, can reveal activation patterns. Here, color indicates first activation of ROIs, from beginning (green) to end (red) of synchronous event. B) The first synchronous event spread through all of the detected ROIs. C) The second and third (D) synchronous event occured in successively smaller subsets of the detected ROIs. All three events exhibited different activation sequences, and ROIs had substantially different activation time points despite being spatially close to one another. Highlighted time-spans in the attached fluorescence profiles (B, C, D, right) indicate affected period of synchronicity.

### Fluorescence imaging and data analysis

In order to assess the algorithms, *in vivo* recorded Ca^2+^ signals of three transgenic mice expressing GCaMP3 were analyzed. Thereby, the animals were imaged first during anesthesia and subsequently while being awake. Three to four FOVs per animal were recorded in the somatosensory cortex, resulting in a total of 23 image sequences. Similar studies, investigating the effects of common anesthetics as well as natural sleep on astroglial Ca^2+^ signaling in the neocortex had previously been performed (17, 18, 25) and served as reference.

First, Ca^2+^ events were analyzed in a generalized context, not differentiating between somatic events and events in the gliapil. In addition, an intersected analysis was performed. Therefore, cell somata were marked manually and intersected with the automatically detected ROIs, resulting in two distinct ROI sets.

Next, the algorithms of MSparkles were compared to three other applications for Ca^2+^ analysis, CHIPS (22), CaSCaDe (7) and AQuA (9). For this comparison, one FOV was randomly selected per mouse, and analyzed in both conditions.

Finally, to demonstrate the wide applicability of PBasE, CoRoDe as well as the analysis capabilities of MSparkles, neuronal Na^+^ signals obtained in acute hippocampal slices were analyzed and compared to current literature. For the analysis of ratiometric Na^+^ activity, ROIs were manually created in ImageJ and imported.

### Adaptive *F*_0_ estimation is able to preserve slow and long-lasting elevations of fluorescence levels

The PBasE *F*_0_ estimation algorithm presented here provides two methods for signal cleanup, a temporal mean filter and a Hampel filter, in order to adapt to different scenarios. In the presence of moderate fluctuations in basal fluorescence, the temporal mean filter and the Hampel filter both produce similar results (Figure 1 A). In the presence of slow, but strong increases of fluorescence levels (Figure 1 B), the Hampel filter typically incorporates these increases into the background, whereas the temporal mean filter is able to preserve such long lasting and slowly rising transients (Figure 1 B).

Pixels covered by the *F*_0_-mask (Figure 1 E) were excluded from the *F*_0_ estimation and set to their respective pre-processed time-course. This results in 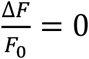 for the affected pixels and thus prevents the detection of false ROIs and subsequently false transients. A side effect of this approach allows to gain a linear speedup of the *F*_0_ computation, corresponding to the percentage of excluded pixels.

### Accurate event detection using the CoRoDe algorithm

ROIs generated by the CoRoDe algorithm were compared against a matching set of 41 manually curated ground-truth ROIs, as well as ROIs obtained using a watershed transform. For the CoRoDe algorithm as well as the watershed transform the range projection *R* was thresholded at 0.6 Δ*F*/*F*_0_. Ground-truth ROIs were carefully outlined using ImageJ, such that the largest visible extent of a fluorescence event was captured. Areas of ground-truth ROIs were compared to the areas of the detected ROIs by computing mean differences as well as relative and absolute size differences (Table 1). Integrated ROI traces (Figure 2 C) were assessed by computing the mean signal-to-noise-ratio (SNR) over all detected ROIs. Since 0.5 Δ*F*/*F*_0_ was chosen as the lowest boundary for transient classification, all values of a given ROI trace below 0.5 were considered noise.

**Table 1:** Validation of detected ROIs. ROIs detected using CoRoDe are more accurate, than those detected using a watershed transform with identical range threshold.

| | Ground truth | Watershed | $T_{\text{CORR}} = 0.5$ | $T_{\text{CORR}} = 0.25$ |
| --- | --- | --- | --- | --- |
| Mean diff | 0 | 51.2297 | -21.3094 | 4.2631 |
| Mean abs diff | 0 | 51.5689 | 21.8059 | 13.8066 |
| Mean relative diff | 0 | 76.7984 | 29.7339 | 18.9355 |
| Mean areA ($\mu\text{m}^2$) | 59.3139 | 110.5436 | 38.0045 | 63.5769 |
| Signal-to-noise-ratio (db) | 15.6 | 13.9 | 17.2 | 15.7 |

Absolute differences in ROI area with respect to the ground truth were significantly reduced for ROIs detected by CoRoDe, when compared to ROIs obtained by a watershed transform (Figure 2 B, D, Table 1). Further, supposing an appropriate correlation threshold, here 0.25 Δ*F*/*F*_0_, ROIs obtained by the CoRoDe algorithm were found to resemble ground truth ROIs much more accurately, compared to regions obtained by watershed transform (Figure 2 B). ROIs obtained using a watershed transform were not only found to be oversized (Table 1, Figure 2 A, B, bottom), but resulting transients, were suppressed and in some cases missed (Figure 2 C, bottom). The quality of integrated ROI time profiles was assessed by comparing their SNR (Figure 2 E, Table 1). Time profiles, of ROIs obtained using the CoRoDe algorithm showed no difference in SNR with respect to ground truth (Figure 2 E, Table 1). SNR is significantly reduced with time profiles of ROIs obtained using watershed transform (Figure 2 E, Table 1**Fehler! Verweisquelle konnte nicht gefunden werden**.).

### Higher Ca^2+^ fluctuations in awake mice

In awake GCaMP3 reporter mice compared to anesthetized animals an increase in both, number of detected ROIs as well as median ROI area was detected (Table 2, Figure 3 A, B). ROIs detected in anesthetized animals were exclusively located in the gliapil. The minimal number of detected active regions was three times higher in awake animals (Table 2). In conjunction to a 50% increase of median ROI count during wakefulness, the median number of transients detected per ROI also doubled (Table 2, Figure 3 C), resulting in the absolute number of detected events to almost double during wakefulness (Table 2, Figure 3 J).

**Table 2:** Statistical analysis of extracted ROI and transient properties. ROI counts, area, transient count, mean frequency, peak amplitude and transient duration are shown with their respective minima, 25%iles, 50%iles (median), 75%iles and maxima.

|  | Anesthetized |  |  |  |  |  | Awake |  |  |  |  |  |
| --- | --- | --- | --- | --- | --- | --- | --- | --- | --- | --- | --- | --- |
|  | #ROIs | Area (μm<sup>2</sup>) | #Tran | Freq. (Hz) | Amp | Dur | #ROIs | Area (μm<sup>2</sup>) | #Tran | Freq. (Hz) | Amp | Dur |
| Min | 19 | 12.91 | 1 | 0.008 | 0.56 | 2.71 | 64 | 12.17 | 1 | 0.008 | 0.57 | 2.85 |
| 25% | 45 | 16.39 | 1 | 0.019 | 0.67 | 3.19 | 114 | 19.37 | 1 | 0.018 | 0.65 | 3.60 |
| 50% | 80 | 17.50 | 1 | 0.028 | 0.79 | 3.88 | 121 | 30.79 | 1.5 | 0.020 | 0.78 | 4.28 |
| 75% | 116 | 18.74 | 1 | 0.047 | 0.87 | 4.27 | 129 | 150.9 | 2 | 0.046 | 0.99 | 5.96 |
| Max | 168 | 25.08 | 2 | 0.063 | 0.91 | 4.84 | 168 | 208.6 | 4 | 0.063 | 1.85 | 22.51 |

The transient frequency per ROI was only computed, if a ROI contained more than one transient. Further, the frequency per ROI was computed as the average frequency of transients between the first and the last occurrence of a transient and not as average of total number of transients divided by the recorded timespan. Investigating median transient frequencies per ROI, contrary to (25) showed no difference (0.0285 Hz and 0.0203 Hz) (Table 2, Figure 3 D) in anesthetized and awake animals, respectively.

Investigation of transient kinetics (Table 2 Figure 3 E, F, G, H) showed no prolongation of median transient duration during wakefulness, compared to anesthetized animals (Figure 3 F). Although signaling activity was increased (Figure 3 A, C, G, H, Table 1) during wakefulness and stronger fluctuations in fluorescence were detectable in awake animals (Figure 3 E, H) no change in overall median peak amplitude was found (Table 1, Figure 3 E). For further investigation, transients were classified, based on their peak amplitude (Figure 3 H), and assigned into one of three classes, with the intervals [0.5, 1), [1, 1.5), [1.5, ∞), respectively. This analysis revealed no difference in median amplitude among transients in the lower two classes obtained from anesthetized and awake animals. On the other hand, a significant difference was found in the class of strongest amplitudes, were awake animals showed a larger median peak amplitude. Further, based on this classification, the signal composition was computed (Figure 3 J, Table 3). While in all transient classes the absolute number of detected transients increased, a shift in relative frequencies was revealed. In awake animals, the percentage of transients > 1.5 Δ*F*/*F*_0_ amplitude transients decreased. nearly doubled, whereas the percentage of low In awake mice, somatic activity could be observed in 75% of the recorded FOVs, whereas no somatic activity was recorded during anesthesia. In order to analyze somatic events independent of events in the gliapil, an additional ROI set containing manually marked somatic regions was generated and subtracted from the ROIs obtained using the CoRoDe algorithm, resulting in two distinct ROI sets, covering somata and the gliapil, respectively. Comparing somatic events to events occurring in the gliapil (Table 4, Figure 3 K, L, M) revealed similar ranges of both, peak amplitude as well as transient durations in both regions. The obtained transient durations were also comparable between somatic regions and gliapil (Table 4, Figure 3 K). Somatic transients exhibited a higher median peak compared to gliapil transients (Table 4, Figure 3 L). Analyzing raw transient counts revealed about 13 times more transients in the gliapil, compared to somatic regions (Table 4, Figure 3 M). Moreover, two thirds of somatic transients exhibited a peak amplitude of 1.5 Δ*F*/*F*_0_ or greater, whereas half of the transients located in the gliapil had a peak amplitude < 1.0 Δ*F*/*F*_0_ (Figure 3 M).

**Table 3:** Detailed statistical analysis of classified peak amplitudes. Minimal and maximal values as well as 25%iles, median values and 75%iles are compared across both conditions along with the respective number of occurrences.

|  | Anesthetized |  |  | Awake |  |  |
| --- | --- | --- | --- | --- | --- | --- |
| | [0.5, 1.0) | [1.0, 1.5) | $> 1.5$ | [0.5, 1.0) | [1.0, 1.5) | $> 1.5$ |
| **Min** | 0.5873 | 1.169 | 1.524 | 0.5975 | 1.158 | 1.653 |
| **25%ile** | 0.6444 | 1.181 | 1.688 | 0.6607 | 1.176 | 1.745 |
| **Median** | 0.6849 | 1.21 | 1.78 | 0.6958 | 1.213 | 1.993 |
| **75%ile** | 0.6974 | 1.217 | 1.942 | 0.7261 | 1.229 | 2.21 |
| **Max** | 0.7504 | 1.388 | 2.304 | 0.7905 | 1.319 | 2.752 |
| **Count** | 1166 | 308 | 156 | 1949 | 635 | 603 |

**Table 4:** Distinct analysis of Ca^2+^ events occurring in somata and the gliapil. Transient amplitude and duration are compared along minimal and maximal values as well as 25%iles, median values and 75%iles are compared across both conditions besides the respective number of occurrences.

|  | Soma |  |  |  |  | Gliapil |  |  |  |  |
| --- | --- | --- | --- | --- | --- | --- | --- | --- | --- | --- |
|  | Amplitude | Duration | [0.5, 1.0) | [1.0, 1.5) | > 1.5 | Amplitude | Duration | [0.5, 1.0) | [1.0, 1.5) | > 1.5 |
| **Min** | 0.044 | 1.633 | 0.505 | 1.027 | 1.506 | 0.500 | 1.148 | 0.500 | 1.001 | 1.509 |
| **25%ile** | 1.388 | 3.570 | 0.549 | 1.061 | 2.333 | 0.733 | 3.627 | 0.563 | 1.084 | 1.763 |
| **Median** | 2.444 | 4.397 | 0.614 | 1.214 | 2.888 | 1.096 | 5.460 | 0.644 | 1.212 | 2.173 |
| **75%ile** | 3.450 | 8.825 | 0.767 | 1.396 | 3.647 | 1.755 | 10.530 | 0.782 | 1.345 | 2.826 |
| **Max** | 6.317 | 42.880 | 0.891 | 1.499 | 5.685 | 5.738 | 41.430 | 0.999 | 1.498 | 8.302 |
| **Count** | 167 | 167 | 29 | 26 | 112 | 2264 | 2264 | 1262 | 450 | 552 |

### Synchronous events are highly diverse

Ca^2+^ activity within the recorded FOV was considered synchronous, if at least half of the detected ROIs were simultaneously active (Figure 4 A, bottom, dashed line). During anesthesia, no significant synchronous activity was detectable (Suppl. Table 10). In fact, no more than 20% of the detected ROIs were simultaneously active (Suppl. Table 10). During wakefulness, synchronous activity was detectable with synchronicity values exceeding 90% (Suppl. Table 10). Investigating the activation sequences of three consecutive synchronous events revealed three points: (I) Synchronous activity started from a few ROIs and then spread across the field of view (Figure 4 B, C, D). (II) It was neither possible to identify a predominant direction of propagation nor a repetitive propagation pattern (Figure 4). In contrast, some regions directly adjacent to the origin of the synchronous event did not show any considerable fluorescence activity until the very end of the synchronous period (Figure 4 C). (III) During three consecutive synchronous events, there was a considerable overlap in the active ROIs, but they were never 100% identical. Moreover, the number of ROIs participating in a synchronous event degraded in consecutive events (Figure 4 B, C, D). Further, the order of activation was different in consecutive synchronous events (Figure 4, B, C, D). These are only observations from a very limited number of datasets and require further investigation to obtain conclusive results.

### Analysis of Na^+^ signals

Recurrent network Na^+^ oscillations in CA1 pyramidal neurons, generated by disinhibition of the hippocampal network, were reliably analyzed by the algorithms of MSparkles (N=4 slice preparations from three different animals) (Suppl. figure 3). For this analysis, PBasE was used to remove trends in the ratiometric data, by computing (*F* − *F*_0_). These trends were caused by accumulating phototoxic effects in one of the fluorescence channels. MSparkles’ signal analysis algorithm identified the onset of Na^+^ oscillations after wash-in of the saline containing 0 Mg^2+^/bicuculline and reported that the network essentially immediately gained a high level of synchronicity (close to 1) between all analyzed CA1 pyramidal neurons in the field of view (30-40 in a given preparation; Suppl. Figure 3 A-C). Individual transients obtained from neuronal cell bodies were categorized into three groups, namely >5, >7.5 and >10% (corresponding to a change in 4.93, 7.39 and 9.85 mM Na^+^) (Suppl. Figure 3 B). The corresponding heatmap illustrates that transients detected in individual neurons fell into all three groups, but that individual network events generally exhibited either larger (7.5 and 10%) or smaller (5 and 7.5%) peak amplitudes in the contributing cells, respectively (Suppl. Figure 3 E). Peak amplitudes and durations of individual transients were comparable to results published previously (Suppl. Figure 3 F, (26)). There was a weak positive linear correlation (R=0.36) between the peak amplitude and the overall duration of individual Na^+^ transients.

### Comparison with state-of-the-art Ca^2+^ analysis software

ROIs obtained by CoRoDe as well as transients analysed by MSparkles were compared to data generated by three other Ca^2+^ analysis applications, CHIPS (22), CaSCaDe (7), and AQuA (9). To obtain objective and reliable results, the official guidelines and tutorials of each application were used and parameters were optimized. MSparkles and most of the tested applications are equipped to automatically remove false-positive ROIs. Therefore, only the number of false negatives (i.e. not detected events) was assessed.

Although AQuA and MSparkles are able to compute a much larger set of parameters, focus was set on transient kinetics (duration and peak amplitude), and the number of detected ROIs and peaks to maintain comparability. For simplicity, dynamic events detected by AQuA were considered as ROIs.

### Detected kinetics are diverse among analysis tools

To compare the performance of different analysis tools (CHIPS (22), CaSCaDe (7) and AQuA (9)), three FOVs recorded in animals expressing GCaMP3 were chosen randomly. All three FOVs in this comparison exhibit seemingly little Ca^2+^ activity during anesthesia. In the second dataset a Ca^2+^ wave of a single astrocyte was recorded in the awake state. The third dataset contains a large Ca^2+^ wave across the entire FOV during wakefulness.

Comparing ROIs detected by different Ca^2+^ analysis applications revealed differences in ROI shape, size, smoothness and location (Suppl. figure 4, Suppl. Table 9). Moreover, ROI segmentation greatly differs among applications (Suppl. figure 4). For example, ROIs generated by CHIPS appeared more rounded and blob-like, whereas ROIs generated by AQuA appeared rough and fragmented. Moreover, CHIPS appeared to generate larger ROIs, whereas CaSCaDe, AQuA and MSparkles provided finer segmentations for comparable regions.

Overall, all applications detected a significant increase in median amplitude in awake state, compared to anesthesia (Figure 5 A). Looking at individual datasets, the determined transient peak amplitudes across the different applications appeared significantly different between anesthetized and awake states, and in some cases ambiguous (Figure 5 A, Suppl. Table 2, 3, 6). In the first dataset CaSCaDe and AQuA found a decrease in median peak amplitude in awake state, CHIPS detected no significant difference and MSparkles detected an increase in median peak amplitude. A similar picture was shown with the second dataset. Unanimity between all tested applications was only given with the third dataset, exhibiting a large Ca^2+^ wave. However, the magnitude of the increase in median amplitude is, again, ambiguous (Figure 5 A).

**Figure 5:**
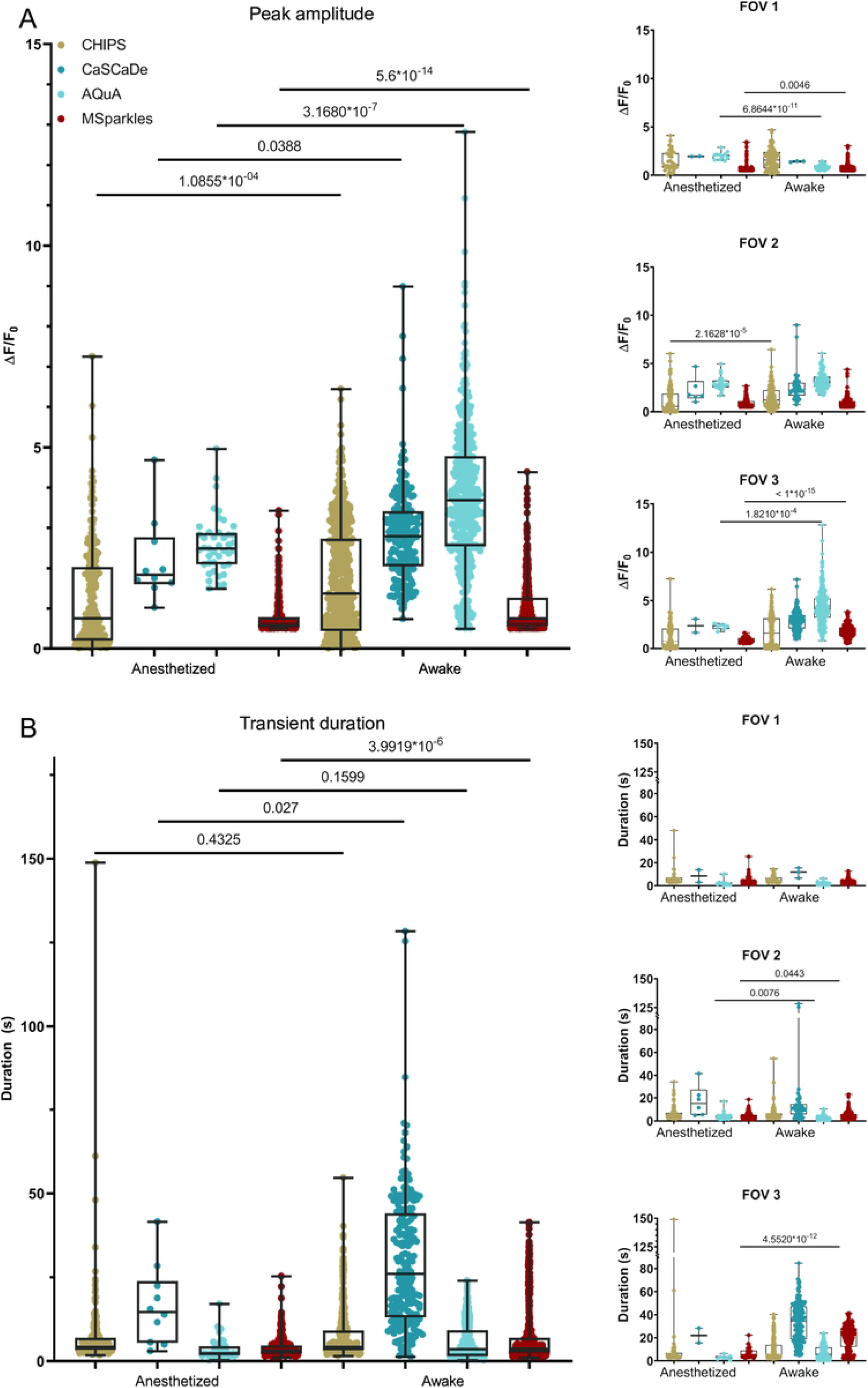
Signal kinetics obtained with different Ca^2+^ analysis tools. A) Obtained peak amplitudes are diverse across different applications and may result in ambiguous tendencies for individual datasets between applications. B) Transient durations are more consistent, however, CaSCaDe tends to measure longer durations than other applications.

Analyzing the transient durations gave similar results (Figure 5 B, Suppl. Table 4, 5, 7). Overall, all applications detected an increased median transient duration in awake animals. However, looking at individual datasets, only AQuA and MSparkles were able to reported significant differences between corresponding anesthetized and awake datasets (Figure 5 B). Interestingly, the transient durations reported by CaSCaDe were 2x – 3x longer, compared to the other applications.

### Time profiles of integrated ROIs show different signatures

In order to investigate the origin of differences in the transient peaks and durations, individual time profiles of ROIs were inspected (Figure 6 A). Therefore, ROIs with a high resemblance across all applications were carefully selected. The detailed investigation of these ROIs in combination their corresponding time profiles revealed several differences. (I) Noise levels of the integrated time profiles varied across the applications (Figure 6 A, right). (II) CHIPS resolved ROIs more coarsely and could not detrend integrated fluorescence profiles. (III) CaSCaDe appeared to prefer extracting transients with a longer duration (Figure 6 A). (IV) Integrated time profiles of AQuA and MSparkles were similar, though the profiles extracted by MSparkles appeared smoother (Figure 6 A, B).

**Figure 6:**
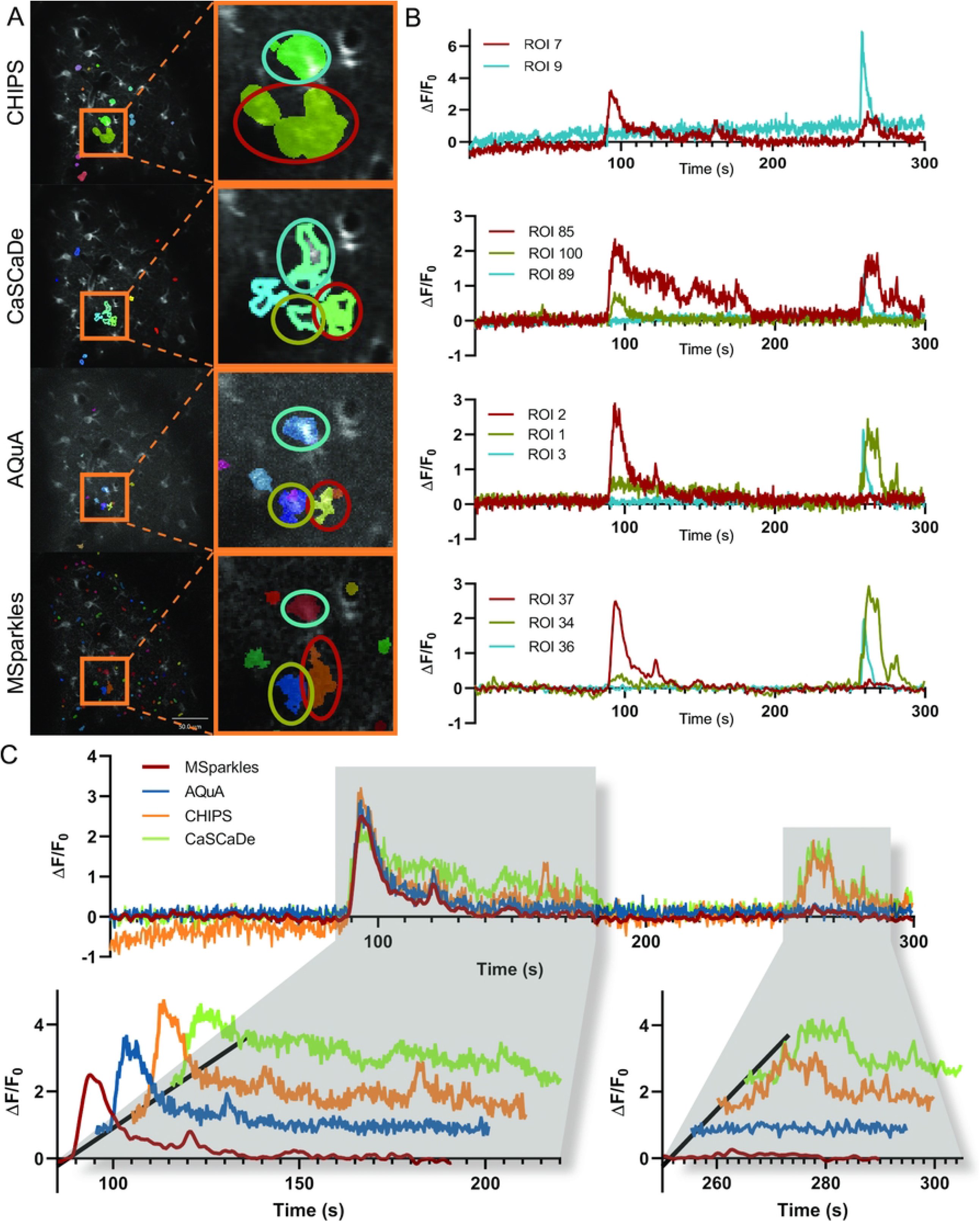
Comparison of ROI detectors. A) ROIs detected with by individual Ca^2+^ analysis tools. Highlighted areas contain fluorescence activity, similarly detected by all tested applications. Comparing ROIs of the magnified areas reveals segmentation differences, as well as differences in size among Ca^2+^ analysis tools. B) Fluorescence profiles of magnified ROIs marked with red, yellow and blue ellipsesPprofiles obtained using CHIPS show an increase in background fluorescence over time. Especially the profiles obtained with CHIPS and CaSCaDe differ to those obtained with AQUA and MSparkles. AQuA and MSparkles performed a more accurate segmentation between the ROIs marked with red and yellow ellipses, which is reflected by the corresponding fluorescence profiles. C) Direct comparison of fluorescence profiles marked by the red ellipse. Profiles are similar around the first event occurring between 90s and 180s. Profiles by CHIPS and CaSCaDe show a third peak and prolonged event, respectively. CHIPS and CaSCaDe show a second fluorescence event at around 270s. AQuA and MSparkles detected this as a separate event, located at the ROI highlighted in yellow.

In addition, the integrated time profiles of a single selected ROI, nearly identically found by all applications were overlaid directly (Figure 6 B). All applications extracted a profile with high resemblance during the first 200 seconds. Especially the prominent transient peak occurring after about 100 seconds is mostly identical across applications. Time profiles integrated by CHIPS and CaSCaDe contained a secondary prominent peak near the end of the recording.

## Discussion

### *F*_0_-estimation with PBasE

Accurate estimation of fluorescence levels at basal concentrations (*F*_0_) of Ca^2+^ and other important messenger molecules is crucial to extract low amplitude transients such as microdomain events, especially in the gliapil (17). During *in-vivo* imaging, in particular during long-term imaging, basal fluorescence levels can vary. These variations do not necessarily occur homogeneously throughout the field of view, requiring pixel precise estimates of *F*_0_. Fitting a low-order polynomial curve to a signal before and after the occurrence of a transient shows accurate results (24). Recently, an adaptive algorithm to automatically estimate *F*_0_ was introduced and verified by comparing it to a reference signal, recorded in a secondary fluorescence channel (17). Approaches, based on biophysical principles (17, 24) allow to reveal the time profile of fluorescence changes (17), and make it possible to detect low amplitude events close to noise level. Similar to the algorithm presented in (24), the PBasE algorithm performs polynomial fitting to estimate fluorescence levels at basal Ca^2+^ concentrations. In addition, it is automated and provides two statistics-based methods to automatically exclude fluorescence transient from baseline estimation. The Hampel filter allows to closely follow slow fluctuations of given signal and is able to exclude relatively short peaks (Figure 1 A). Thereby, slow and long-lasting fluorescence elevations are incorporated into the estimate baseline. This may be desirable to compensate for slowly rising basal fluorescence and closely resembles the behavior presented in (17). The temporal mean filter on the other hand is capable to preserve such slow and long-lasting elevations for later analysis. Automatic signal stabilization not only prevents high frequency oscillations, but makes this algorithm suitable for long-term recordings.

### Automated ROI detection

In combination with PBasE the CoRoDe algorithm was able to detect a plethora of ROIs containing low amplitude events, like microdomain events, not easily visible to a human observer and also not detectable by most other Ca^2+^ analysis applications. These ROIs were predominantly located in the gliapil, where the majority of Ca^2+^ events occur (14). The computed *F*_0_ baseline permits effectively removing slow fluctuations in background fluorescence and thus enables the extraction of subtle events, otherwise obscured by these fluctuations. In addition, the CoRoDe algorithm is capable of extracting active regions more precisely than the commonly used watershed segmentation. This in turn permits subsequent ROI integration to accurately extract transients. Using the range *R* of Δ*F*/*F*_0_ for ROI extraction has the advantage to only project actual changes in Δ*F*/*F*_0_, in contrast to using maximum or summed intensity projections which do not necessarily correspond to fluorescence events and tend to suppress weak events. However, if multiple events overlap during the recorded time period, these events might not be resolved properly and in some cases might not be detected. This is a general shortcoming of projection-based ROI detectors.

### Analysis of Ca^2+^ transients

Analyzing Ca^2+^ transients in awake and anesthetized mice showed increased ROI numbers (Figure 3 A,), ROI area (Figure 3 B) as well as transients per ROI (Figure 3 C) in awake animals. Furthermore, awake animals exhibited about twice as many Ca^2+^ transients (Figure 3 J) as well as higher peak amplitudes (Figure 3 E, G). This overall increased Ca^2+^ activity in awake animals is in line with previous studies (17, 18, 25). No significant decrease in the duration of Ca^2+^ transients was detected in awake animals (Figure 3 F). Instead, some animals displayed a prolonged transient duration. This might be attributable to the presence of large and long lasting Ca^2+^ waves, which occurred exclusively in awake animals and were detected by MSparkles as individual events.

Classifying Ca^2+^ transients based on their peak amplitude allows to calculate the signal-composition, i.e. the relative frequency of amplitude peaks within a defined interval. This facilitates the detection of changes in the relative incidence of the respective transient classes. Changes in signal composition can provide a more meaningful statement, than just median peak amplitude. Visualizing the signal composition shows not only the overall reduction of Ca^2+^ activity, but further illustrates that this reduction happens largely on the cost of high amplitude transients, which is line with previous work (17, 25).

### Synchronous Ca^2+^ events occur during wakefulness

Synchronous Ca^2+^ event activity was defined at a threshold of 50% simultaneously active ROIs. Synchronous activity, was detected exclusively during wakefulness, as observed by others (25). Most of the synchronous activity can directly be attributed to animal motion. In addition, detailed investigation of multiple, consecutive synchronous events showed that these events are highly diverse. Each of the detected events showed a different activation order, and no detectable activation pattern. However, with each reoccurring synchronous event, the number of participating astrocytes decreased.

### Comparison with other software

The properties of the algorithms PBasE and CoRoDe, implemented in MSparkles, were compared to three published Ca^2+^ signal analysis applications. All of the tested applications detected similar overall trends (Figure 5 A, B), although the mean values and value ranges differed substantially among applications. To our surprise, the individual results per FOV were quite diverse (Figure 5 A, B, Suppl. Table 2-9). Not only did the number of detected ROIs differ substantially, but also the number of extracted transients and more importantly the resulting individual trends. Investigating ROIs with high resemblance across the tested applications in combination with their corresponding time profiles provided a possible answer for this diversity. During our evaluation, CHIPS had problems in background correction. This not only has an effect on the scaling of signal transients when computing Δ*F*/*F*_0_ but can further cause problems and result in inaccurate results when detecting and analyzing transient amplitudes (Figure 6 A, right). The ROIs subjected to further investigation integrated with CHIPS and CaSCaDe contained a secondary prominent transient peak (Figure 6 A, B). Careful analysis revealed this peak originating from a single ROI that had been segmented into two distinct ROIs by the other analysis applications. Only AQuA and MSparkles were able to resolve these ROIs properly. Despite AQuA performing dynamic event analysis and MSparkles performing a classical ROI analysis, the compared time profiles are very similar, with the difference being that profiles extracted by MSparkles’ appear smoother (Figure 6 A, B).

Transient durations computed with CaSCaDe were two to three times longer than reported by any other application. Transient peaks reported by AQuA tend to exhibit a higher amplitude compared to any other analysis. However, AQuA reports fluorescence values based on local maxima, in contrast to averaged values reported by the other applications.

AQuA and MSparkles provide a full graphical user interface, granting also non-programming experts access to advanced fluorescence analysis. MSparkles further provides real-time visual feedback and interactive previews for data exploration and algorithm optimization. As an effort to work towards a common standard within the Ca^2+^ analysis community we provide extensive definitions and explanations of our terminology and computed transient properties (Suppl. Table 1 Suppl. figure 2).

## Conclusion

To quantitatively evaluate fluorescent signal recordings of brain tissues with varying signal-to-noise ratios, vastly different fluorescence levels, recording artefacts and diverse temporal resolutions, two novel algorithms were developed. PBasE for adaptive *F*_0_estimation and CoRoDe for the detection of fluorescently active stationary regions. These algorithms made it possible to identify a large number of ROIs at high spatial resolution from which close-to-noise signal (Ca^2+^ or Na^+^) transients could be determined. The analysis algorithms are embedded in a graphical user interface MSparkles which assists in data analysis without programming skills.

## Materials and Methods

### Cranial window surgery for *in vivo* two-photon imaging

During surgical procedures, animals were kept on heating pads and eyes were covered with Bepanthen ointment (Bayer). Anesthesia was induced with a mixture of 5 % isoflurane, 47.5 % O_2_ (0.6 l/ min) and 47.5 % N_2_O (0.4 l/ min) and maintained with 2 % isoflurane (Harvard Apparatus anesthetic vaporizer). A standard craniotomy (27) of 3 mm in diameter was performed over the somatosensory cortex (2 mm posterior and 1.5 mm lateral to bregma). The craniotomy was sealed with a glass coverslip and fixed with dental cement (RelyX®, 3M ESPE). Subsequently, a custom-made metal holder for head restraining (5 mm diameter) was applied and fixed to the skull with dental cement. After surgery, the animals were kept on the heating pad until complete recovery and received pain treatment for at least three days (carprofen/burprenorphine). After five to seven days the first imaging session was performed.

### Two-photon laser scanning microscopy (2P-LSM)

*In vivo* 2P-LSM was performed using a custom-built microscope equipped with a resonant scanner (RESSCAN-MOM, Sutter instrument) and a 20x water-immersion objective (W Plan-Apochromat 20x/1.0 DIC D=0.17 M27; Zeiss). Images were acquired with a frame rate of 30 Hz and a frame-averaging factor of 10, resulting in an effective acquisition rate of 3 Hz. Recorded fields of view (FOVs) had a size of 256 × 256 µm, sampled with 512 × 512 pixels (0.5 µm/pixel). To minimize photo-damage, incident laser power was kept between 30 – 40 mW for a sufficient signal-to-noise ratio. Laser wavelength was set to 890 nm (Chameleon Ultra II, Ti:Sapphire Laser, Coherent). The emitted light was detected by photomultiplier tubes (R6357, Hamamatsu) (27) and pre-amplified (DHPCA-100, Femto). Digitizer (NI-5734) and control hardware (NI-6341) were housed in a NI PXIe (1082) chassis, connected to a control-PC via a high bandwidth PXIe-PCIe8398 interface. Scanning and image acquisition were controlled by ScanImage (SI 5.6R1) (28).

### *In vivo* two-photon Ca^2+^ imaging

In preparation for Ca^2+^ imaging, animals were habituated before the first imaging session according to adapted protocols without water restriction (29, 30). The animals were head-fixed with a custom-designed head-restrainer, 3D-printed using stainless steel. During imaging, anesthesia was applied using a custom-made, magnetically attachable mask. Each FOV was imaged twice: first in anesthetized, then in awake state. During imaging in anesthetized state, isoflurane concentration was kept at 1.5 %, and flow of O_2_ and N_2_O was set to 0.6 l/min and 0.4l /min, respectively. Before awake state imaging, isoflurane and other gases were switched off and it was visually verified that animals were fully awake (mice were grooming, voluntarily moving or reacting to air puffs). The selected FOVs for Ca^2+^ imaging were located in the somatosensory cortex, 80 – 100 μm beneath the dura. Each FOV was recorded for 5 min per condition to investigate Ca^2+^ events. The total duration of one imaging session ranged between 30-60 min per animal. Anesthetized mice were kept on a heating pad at 37°C until they recovered completely. In addition, mice had access to high-caloric food (Fresubin, Fresenius Kabi GmbH) *ad libitum*.

### Wide-field *in situ* Na^+^ imaging

*Balb/c* mice aged between postnatal day (P) 14-18 were anesthetized with CO_2_ before being quickly decapitated and 250 µm hippocampal slices were prepared as previously described (31, 32). As reported in detail in (31, 32), slices were transferred into an experimental bath, continuously perfused with standard, CO_2_/HCO_3-_-buffered artificial cerebrospinal fluid (ACSF) and their CA1 region bolus-stained with the Na^+^ sensitive dye SBFI-AM (sodium-binding benzofuran isophthalate-acetoxymethyl ester; Invitrogen, Karlsruhe, Germany). SBFI was alternatively excited at 340/380 nm at an imaging frequency of 0.5 Hz and emission was collected >440 nm from defined regions of interest (ROIs) reflecting cell bodies of CA1 pyramidal neurons. Changes in the SBFI ratio were transferred into changes in intracellular Na^+^ concentration based on *in situ* calibrations (31, 32). Recurrent network activity was induced via perfusion with an ACSF lacking Mg^2+^ and containing 10 µM bicuculline, in order to remove the Mg^2+^ block from NMDA receptors, and to prevent GABA_A_ receptor activation respectively (32).

### Animals

Astrocyte-specific knockin *GLAST-CreERT2* mice (*Slc1a3tm1(cre/ERT2)Mgoe, MGI:3830051*) (33) were crossbred to Rosa26 reporter mice with GCaMP3 expression (*Gt(ROSA)26Sortm1(CAG-GCaMP3)Dbe, MGI: 5659933*) (34). Imaging sessions were performed at 8-10 weeks of age, 21 days after tamoxifen induced recombination (35). GCaMP3 expression was induced in *Glast*-*CreERT2* mice, tamoxifen (Carbolution, Neunkirchen, Germany) was intraperitoneally injected (100 μL/10 g body weight with 10 μg/mL Tamoxifen in Mygliol®812 (Caesar and Lorentz GmbH, Hilden, Germany)) to mice once per day for three consecutive days.

### Statistical analysis and figures

Statistical analysis of computed data was conducted using GraphPad Prism 8. D’Agostino-Pearson normality test was used to test for were normally or log-normally distributed values. For non-normal distributions, median values, with their associated ranges, inter quartile ranges and percentiles were used for statistical evaluation. Similarly, statistical significances were computed using the Mann-Whitney test for individual, or Kruskal-Wallis test for multiple comparisons, as non-parametric tests, suitable for non-normal distributions.

Figures were arranged using Adobe InDesign 2020, Adobe Illustrator 2020 and GraphPad Prism 8. Graphs, trace plots, kymographs, and ROI maps were directly exported from MSparkles. Additional ROI maps were extracted from the respective Ca^2+^ analysis tools.

### Pre-processing

Pre-processing was adjusted individually per dataset as needed using MSparkles built-in pre-processing pipeline (Suppl. figure 1 B). Independent of individual settings, all datasets were denoised (SURE-LET (36)) and subjected to a temporal median filter. No external software was used. The preprocessed dataset is referred to as *F*.

## Code availability

https://gitlab.com/Gebhard/MSparkles/

## Data availability

Imaging datasets will be made available upon request.

## Author contributions

GS developed and programmed PBase, CoRoDe and MSparkles, analyzed data, wrote first manuscript draft and prepared the figures; LCC, LS, LF and KE performed experiments, analyzed data and tested the software; PR, DG and XB analyzed data and tested the software. CRR, AS and FK supervised the project, provided grant support and revised text and figures. All authors provided critical input at various steps of the project and agreed on the final version of the manuscript.

## Acknowledgments

Carmen Kasakow and Wenhui Huang provided valuable help and support during testing PBase, CoRoDe and MSparkles.

## Funding

This work has been supported by grants from the European Union’s Horizon 2020 FETPROACT-01-2016 Neurofibers, the H2020 MSCA-ITN EU-GliaPhD, from the Deutsche Forschungsgemeinschaft (DFG) SPP 1757 (to CRR, FK), DFG SFB894 (to FK), DFG FOR2289 (to AS, FK) and DFG SFB1158 (to FK).

## Competing interests

We declare no competing interests.

## Ethics statement

For Ca^2+^ imaging experiments conducted in Homburg, Germany, mice were kept and bred in strict accordance with the recommendations to European and German guidelines for the welfare of experimental animals. Animal experiments were approved by the Saarland state’s “Landesamt für Gesundheit und Verbraucherschutz” in Saarbrücken/Germany (veterinary licenses: 71/2013, 36/2016).

Na^+^ imaging experiments conducted at the Heinrich Heine University Düsseldorf, Germany, were carried out in accordance with the institutional guidelines and the European Community Council Directive (86/609/EEC). All experiments were approved by the Animal Welfare Office at the Animal Care and Use Facility of the Heinrich Heine University Düsseldorf (institutional act number: O52/05.

## Supporting information captions

**Supplementary figure 1: User interface and processing pipeline**. A) MSparkles main user interface including hierarchical data management (left) the currently loaded dataset with detected ROIs (center) as well as analyzed and classified time profiles (bottom). Moving the mouse across the loaded dataset, displays the original and the normalized time profiles (top-right) of the pixel currently underneath the mouse pointer. B) The Processing pipeline with individually executable and configurable pipeline stages. Pre-processing itself provides a customizable pipeline, allowing to add and remove algorithms and filters as needed. After F_0_ estimation, one or more ROI detection methods can be configured. Finally, the analysis stage performs ROI integration and other downstream computations. Obtained results and figures and graphs are automatically exported.

**Supplementary table 5: Definition of terminology**. Definition of terminology and quantities used in MSparkles.

**Supplementary figure 2: Properties of Ca**^**2+**^ **transients**. A) Properties of individual transients include peak amplitude (yellow dot), duration (orange lines) as well as rise (green) and decay times (red). Depending on the height reference, 50% (FWHM, orange line), 25%, or 10% (dashed orange lines), the duration, but also rise and decay times can be calculated more accurately. This however, requires a high signal quality. B) Consecutive transients allow to compute various signal timings, like peak-to-peak, inter signal and start-to-start. It is important to notice that these timings are influenced by the choice of the height reference.

**Supplementary figure 3: Evaluation if Na**^**+**^ **signals analyzed with MSparkles**. (A) Image of the CA1 pyramidal cell layer of a hippocampal slice (P16) stained with sodium-binding benzofuran isophthalate-AM (SBFI-AM), scale bar is 25 µm. Circles represent regions 1-5 as depicted in (B). (B) Na^+^ signals from regions 1-5 as detected during recurrent network activity. Peaks detected by the software shown by colored dots depending on threshold groups (Red >10% Green >7.5%, Blue >5%). Peak amplitude and full width at half maximum are indicated by black lines. (C) Synchronicity plot of all cells measured in the experiment shown in (A) and (n=33), showing the proportion of cells with activity over time. (D) 3D plot generated by MSparkles, showing Na^+^ traces of all measured cells. (E) Threshold group heat map showing the time points at which each cell was involved in peaks with color code corresponding to that in (B). (F) Scatter plot generated by MSparkles showing the correlation between the duration and amplitude of signals. This figure and therein displayed results have been generated using MSparkles at the University of Düsseldorf. Figure by Lisa Felix & Katharina Everaerts, HHU Düsseldorf.

**Supplementary figure 4: Detailed comparison of ROI detectors**. Each column shows detected ROIs obtained with a specific Ca^2+^ analysis tool. Each row is dedicated to a single FOV. Highlighted regions point out differences in detector sensitivity as well as region segmentation, potentially resulting in ambiguous measurements of ROI sizes and thus differences in ROI integration and resulting peak amplitudes. CHIPS tends to extract large and smooth regions. CaSCaDe extracts regions with a high degree of segmentation. Regions extracted by AQuA tend to be rough and contain holes. MSparkles is able to extract regions with varying smoothness, based on the temporal correlation of pixels. Due to the interplay of the PBasE and CoRoDe algorithms, MSparkles is able to detect active regions with localized and dim fluorescence events. Scale bar 50 µm.

**Supplementary table 6: P-values between all peak amplitudes by analysis tool during anesthesia**. P-values indicate statistically significant differences between the results obtained by different analysis applications analysing transients of anesthetized mice.

**Supplementary table 7: P-values between all peak amplitudes by analysis tool in awake state**. P-values indicate statistically significant differences between the results obtained by different analysis applications analysing transients of awake mice.

**Supplementary table 8: P-values between all measured signal durations by analysis tool during anesthesia**. P-values indicate statistically significant differences between the results obtained by different analysis applications analysing transients of anesthetized mice.

**Supplementary table 9: P-values between all measured signal durations by analysis tool in awake state**.P-values indicate statistically significant differences between the results obtained by different analysis applications analysing transients of awake mice.

**Supplementary table 10: Comparison of median peak amplitudes**. Median peak amplitude per field of view, extracted by Ca^2+^ analysis applications.

**Supplementary table 11: Comparison of median transient durations**. Transient duration extracted by Ca^2+^ analysis applications.

**Supplementary table 12: Comparison of detected ROIs**. Detected ROIs, false negative ROIs and signal counts. False negatives were assessed by careful manual evaluation in ImageJ for each application.

**Supplementary table 13: Mean ROI areas per FOV as detected by applications**. Mean areas of detected ROIS wich corresponding standard deviation, per FOV and condition.

**Supplementary table 14: Maximum synchronicity of GCaMP3 animals in percent**. FOVs exhibiting a high synchronous activity above 50% are marked green.

## Notes

### Competing Interest Statement

The authors have declared no competing interest.

## References

1. Giaume C, Kirchhoff F, Matute C, Reichenbach A, Verkhratsky A. Glia: the fulcrum of brain diseases. Cell DeathDiffer. 2007;14(7):1324–35.

2. Araque A, Parpura V, Sanzgiri RP, Haydon PG. Tripartite synapses: glia, the unacknowledged partner. Trends Neurosci. 1999;22(5):208–15.

3. Alberdi E, Sánchez-Gómez MV, Matute C. Calcium and glial cell death. Cell Calcium. 2005;38(3-4):417–25.

4. Caudal LC, Gobbo D, Scheller A, Kirchhoff F. The Paradox of Astroglial Ca2 + Signals at the Interface of Excitation and Inhibition. Frontiers in Cellular Neuroscience. 2020;14(399).

5. Ellefsen KL, Settle B, Parker I, Smith IF. An algorithm for automated detection, localization and measurement of local calcium signals from camera-based imaging. Cell Calcium. 2014;56(3):147–56.

6. Cheng H, Song LS, Shirokova N, González A, Lakatta EG, Ríos E, et al. Amplitude distribution of calcium sparks in confocal images: theory and studies with an automatic detection method. Biophys J. 1999;76(2):606–17.

7. Agarwal A, Wu PH, Hughes EG, Fukaya M, Tischfield MA, Langseth AJ, et al. Transient Opening of the Mitochondrial Permeability Transition Pore Induces Microdomain Calcium Transients in Astrocyte Processes. Neuron. 2017;93(3):587-605.e7.

8. Srinivasan R, Huang BS, Venugopal S, Johnston AD, Chai H, Zeng H, et al. Ca(2+) signaling in astrocytes from Ip3r2(-/-) mice in brain slices and during startle responses in vivo. Nat Neurosci. 2015;18(5):708–17.

9. Wang Y, DelRosso NV, Vaidyanathan TV, Cahill MK, Reitman ME, Pittolo S, et al. Accurate quantification of astrocyte and neurotransmitter fluorescence dynamics for single-cell and population-level physiology. Nat Neurosci. 2019;22(11):1936–44.

10. Picht E, Zima AV, Blatter LA, Bers DM. SparkMaster: automated calcium spark analysis with ImageJ. Am J Physiol Cell Physiol. 2007;293(3):C1073–81.

11. Giovannucci A, Friedrich J, Gunn P, Kalfon J, Brown BL, Koay SA, et al. CaImAn an open source tool for scalable calcium imaging data analysis. Elife. 2019;8.

12. Oberheim NA, Goldman SA, Nedergaard M. Heterogeneity of astrocytic form and function. Methods in molecular biology (Clifton, NJ). 2012;814:23–45.

13. Nimmerjahn A, Mukamel EA, Schnitzer MJ. Motor behavior activates Bergmann glial networks. Neuron. 2009;62(3):400–12.

14. Bindocci E, Savtchouk I, Liaudet N, Becker D, Carriero G, Volterra A. Three-dimensional Ca(2+) imaging advances understanding of astrocyte biology. Science. 2017;356(6339).

15. Shigetomi E, Bushong EA, Haustein MD, Tong X, Jackson-Weaver O, Kracun S, et al. Imaging calcium microdomains within entire astrocyte territories and endfeet with GCaMPs expressed using adeno-associated viruses. J Gen Physiol. 2013;141(5):633–47.

16. Smith IF, Parker I. Imaging the quantal substructure of single IP3R channel activity during Ca2+ puffs in intact mammalian cells. Proc Natl Acad Sci U S A. 2009;106(15):6404–9.

17. Mueller FE, Cherkas V, Stopper G, Caudal LC, Stopper L, Kirchhoff F, et al. Deciphering spatio-temporal fluorescence changes using multi-threshold event detection (MTED). bioRxiv. 2020:2020.12.06.413492.

18. Bojarskaite L, Bjørnstad DM, Pettersen KH, Cunen C, Hermansen GH, Åbjørsbråten KS, et al. Astrocytic Ca. Nat Commun. 2020;11(1):3240.

19. MATLAB. version 9.8 (R2020a). The MathWorks Inc. Natick, Massachusetts; 2020.

20. Thrane AS, Thrane VR, Zeppenfeld D, Lou N, Xu Q, Nagelhus EA, et al. General anesthesia selectively disrupts astrocyte calcium signaling in the awake mouse cortex. Proceedings of the National Academy of Sciences of the United States of America. 2012;109(46):18974–9.

21. Bojarskaite L, Bjørnstad DM, Pettersen KH, Cunen C, Hermansen GH, Åbjørsbråten KS, et al. Astrocytic Ca^2+^ signaling is reduced during sleep and is involved in the regulation of slow wave sleep. Nat Commun. 2020;11(1):3240.

22. Barrett MJP, Ferrari KD, Stobart JL, Holub M, Weber B. CHIPS: an Extensible Toolbox for Cellular and Hemodynamic Two-Photon Image Analysis. Neuroinformatics. 2018;16(1):145–7.

23. Hampel FR. The Influence Curve and Its Role in Robust Estimation. Journal of the American Statistical Association. 1974;69(346):383–93.

24. Balkenius A, Johansson AJ, Balkenius C. Comparing Analysis Methods in Functional Calcium Imaging of the Insect Brain. PLoS One. 2015;10(6):e0129614.

25. Thrane AS, Rangroo Thrane V, Zeppenfeld D, Lou N, Xu Q, Nagelhus EA, et al. General anesthesia selectively disrupts astrocyte calcium signaling in the awake mouse cortex. Proc Natl Acad Sci U S A. 2012;109(46):18974–9.

26. Karus C, Mondragao MA, Ziemens D, Rose CR. Astrocytes restrict discharge duration and neuronal sodium loads during recurrent network activity. Glia. 2015;63(6):936–57.

27. Cupido A, Catalin B, Steffens H, Kirchhoff F. Surgical procedures to study microglial motility in the brain and in the spinal cord by in vivo two-photon laser-scanning microcopy. In: Bakota L, Brandt R, editors. Confocal and Multiphoton Laser-Scanning Microscopy of Neuronal Tissue: Applications and Quantitative Image Analysis. 87: Springer; 2014. p. 37–50.

28. Pologruto TA, Sabatini BL, Svoboda K. ScanImage: flexible software for operating laser scanning microscopes. Biomed Eng Online. 2003;2:13.

29. Guo ZV, Hires SA, Li N, O’Connor DH, Komiyama T, Ophir E, et al. Procedures for behavioral experiments in head-fixed mice. PLoS One. 2014;9(2):e88678.

30. Kislin M, Mugantseva E, Molotkov D, Kulesskaya N, Khirug S, Kirilkin I, et al. Flat-floored air-lifted platform: a new method for combining behavior with microscopy or electrophysiology on awake freely moving rodents. J Vis Exp. 2014(88):e51869.

31. Felix L, Ziemens D, Seifert G, Rose CR. Spontaneous Ultraslow Na(+) Fluctuations in the Neonatal Mouse Brain. Cells. 2019;9(1).

32. Karus C, Mondragão MA, Ziemens D, Rose CR. Astrocytes restrict discharge duration and neuronal sodium loads during recurrent network activity. Glia. 2015;63(6):936–57.

33. Mori T, Tanaka K, Buffo A, Wurst W, Kühn R, Götz M. Inducible gene deletion in astroglia and radial glia--a valuable tool for functional and lineage analysis. Glia. 2006;54(1):21–34.

34. Paukert M, Agarwal A, Cha J, Doze VA, Kang JU, Bergles DE. Norepinephrine controls astroglial responsiveness to local circuit activity. Neuron. 2014;82(6):1263–70.

35. Jahn HM, Kasakow CV, Helfer A, Michely J, Verkhratsky A, Maurer HH, et al. Refined protocols of tamoxifen injection for inducible DNA recombination in mouse astroglia. Sci Rep. 2018;8(1):5913.

36. Luisier F, Blu T, Unser M. A new SURE approach to image denoising: interscale orthonormal wavelet thresholding. IEEE Trans Image Process. 2007;16(3):593–606.

